# An Open MRI Dataset for Multiscale Neuroscience

**DOI:** 10.1101/2021.08.04.454795

**Authors:** Jessica Royer, Raúl Rodríguez-Cruces, Shahin Tavakol, Sara Larivière, Peer Herholz, Qiongling Li, Reinder Vos de Wael, Casey Paquola, Oualid Benkarim, Bo-yong Park, Alexander J. Lowe, Daniel Margulies, Jonathan Smallwood, Andrea Bernasconi, Neda Bernasconi, Birgit Frauscher, Boris C. Bernhardt

## Abstract

Multimodal neuroimaging grants a powerful window into the structure and function of the human brain at multiple scales. Recent methodological and conceptual advances have enabled investigations of the interplay between large-scale spatial trends (also referred to as gradients) in brain microstructure and connectivity, offering an integrative framework to study multiscale brain organization. Here, we share a multimodal MRI dataset for Microstructure-Informed Connectomics (MICA-MICs) acquired in 50 healthy adults (23 women; 29.54±5.62 years) who underwent high-resolution T1-weighted MRI, myelin-sensitive quantitative T1 relaxometry, diffusion-weighted MRI, and resting-state functional MRI at 3 Tesla. In addition to raw anonymized MRI data, this release includes brain-wide connectomes derived from i) resting-state functional imaging, ii) diffusion tractography, iii) microstructure covariance analysis, and iv) geodesic cortical distance, gathered across multiple parcellation scales. Alongside, we share large-scale gradients estimated from each modality and parcellation scale. Our dataset will facilitate future research examining the coupling between brain microstructure, connectivity, and macroscale function. MICA-MICs is available on the Canadian Open Neuroscience Platform’s data portal (https://portal.conp.ca).

## Background & Summary

The human brain is a highly interconnected network which can be described at multiple spatial and temporal scales. Neuroimaging, in particular magnetic resonance imaging (MRI), has provided a window into brain structure and function, offering versatile contrasts to assess its multiscale organization ^1^. Multimodal imaging increasingly capitalizes on sequences sensitive to brain microstructure, such as quantitative T1 (qT1) relaxation mapping. This contrast can differentiate highly myelinated regions, with shorter T1 relaxation times, from more lightly myelinated regions showing longer qT1 ^2^. Regional variations in qT1 concord with seminal myeloarchitectonic studies ^3–5^, supporting the potential of these contrasts for *in vivo* microstructural profiling and the study of myeloarchitectonic similarity between areas ^6–9^. These investigations can also be complemented by metrics such as geodesic distance, enabling estimations of cortico-cortical wiring cost emerging from short-range intracortical axon collaterals ^10–13^, the exploration of the anatomical proximity of different brain systems, and the study of cortical topographic organization ^14,15^. In addition, macroscale connectome architecture can be probed using diffusion MRI tractography and resting-state functional connectivity analysis to approximate whole-brain structural and functional networks ^16–18^. Together, these techniques offer key insights into overarching principles of brain organization, from properties of local regions to their embedding within macroscale systems.

Recent methodological and conceptual advances have provided the means to analyse topographic principles of multiscale brain organization. Homogeneity in regional properties can be detected in structural and functional imaging data, at the basis of parcellation-based approaches ^19^. Regional boundaries can be defined with a varying level of granularity from different features, such as morphology ^20,21^, microstructure ^22,23^, connectivity patterns ^24,25^, and combinations of these metrics ^26^. Functional and anatomical relationships between parcels can then be identified, forming the brain’s macroscale network architecture ^27–29^. Complementing techniques highlighting discrete collections of areas through parcellation or decomposing the brain into mesoscale communities, recent work has begun to identify continuous spatial trends – also referred to as gradients – in brain microstructure, connectivity, and function. Gradient identification approaches have described main axes of cortical and subregional organization at the level of resting-state functional connectivity ^14,30–35^, structural connectivity derived from diffusion tractography ^36–39^, similarity of cortical microstructure ^6,7,13,40–42^ and cortical morphology ^41^, as well as molecular and microcircuit properties ^16,43,44^. These approaches have enabled the discovery of a principal gradient of intrinsic functional connectivity differentiating lower-order sensorimotor systems from transmodal systems such as the default-mode network and paralimbic cortices, recapitulating seminal models of the cortical hierarchy formulated in non-human primates ^6,45,46^. By depicting low dimensional axes of cortical organization, gradient approaches enable investigations of systematic changes in structure and function across the brain and are thus particularly suited for studies aiming to bridge different neurobiological axes. For instance, recent work has demonstrated stronger decoupling between principal microstructural and functional gradients in transmodal cortical areas relative to unimodal systems, possibly reflective of the flexible role that transmodal areas play in human cognition ^6^. Relatedly, the principal functional gradient has also been shown to reflect variations in geodesic distance between sensory and transmodal systems, offering a potential macroscale mechanism allowing transmodal networks to support higher cognitive functions decoupled from “the here and now” ^14^. By offering a formal framework for such multimodal comparisons, these findings emphasize the potential of dimensional analyses to obtain novel insights into multiscale brain organization.

Beyond innovations in imaging and analytics, neuroscience has increasingly benefitted from the adoption of open science practices, particularly through open data sharing ^47–49^. In addition to advancing our understanding of brain organization, these repositories have supported increased exchange and collaboration, boosting transparency and reproducibility ^50^. In line with this perspective, this work presents a ready-to-use multimodal MRI dataset for Microstructure-Informed Connectomics (MICA-MICs). MICA-MICs provides connectomes based on i) task-free functional MRI, ii) diffusion tractography, iii) microstructure covariance analysis based on qT1 mapping, and iv) geodesic cortical distance, each built across multiple parcellation schemes and spatial scales. We furthermore provide anonymized raw data adhering to Brain Imaging Data Structure (BIDS) standards ^51^. Processing has been carried out using an open access pipeline (https://micapipe.readthedocs.io/). This resource promises to deepen our understanding of the human brain at multiple scales and augment assessments of generalizability and replicability.

## Methods

### Participants

Data were collected in a sample of 50 healthy volunteers (23 women; 29.54±5.62 years; 47 right-handed) between April 2018 and February 2021. Each participant underwent a single testing session. All participants denied a history of neurological and psychiatric illness. The Ethics Committee of the Montreal Neurological Institute and Hospital approved the study (2018-3469). Written informed consent, including a statement for openly sharing all data in anonymized form, was obtained from all participants. Socio-demographic information included in this release includes participant sex and age at time of scan (in 5-year increments).

### MRI data acquisition

Scans were completed at the Brain Imaging Centre of the Montreal Neurological Institute and Hospital on a 3T Siemens Magnetom Prisma-Fit equipped with a 64-channel head coil. Participants underwent a T1-weighted (T1w) structural scan, followed by multi-shell diffusion-weighted imaging (DWI) and resting-state functional MRI (rs-fMRI). In addition, a pair of spin-echo images was acquired for distortion correction of individual rs-fMRI scans. A second T1w scan was then acquired, followed by qT1 mapping (**Figure 1A**). Total scan time for these acquisitions was approximately 45 minutes.

**Figure 1.**
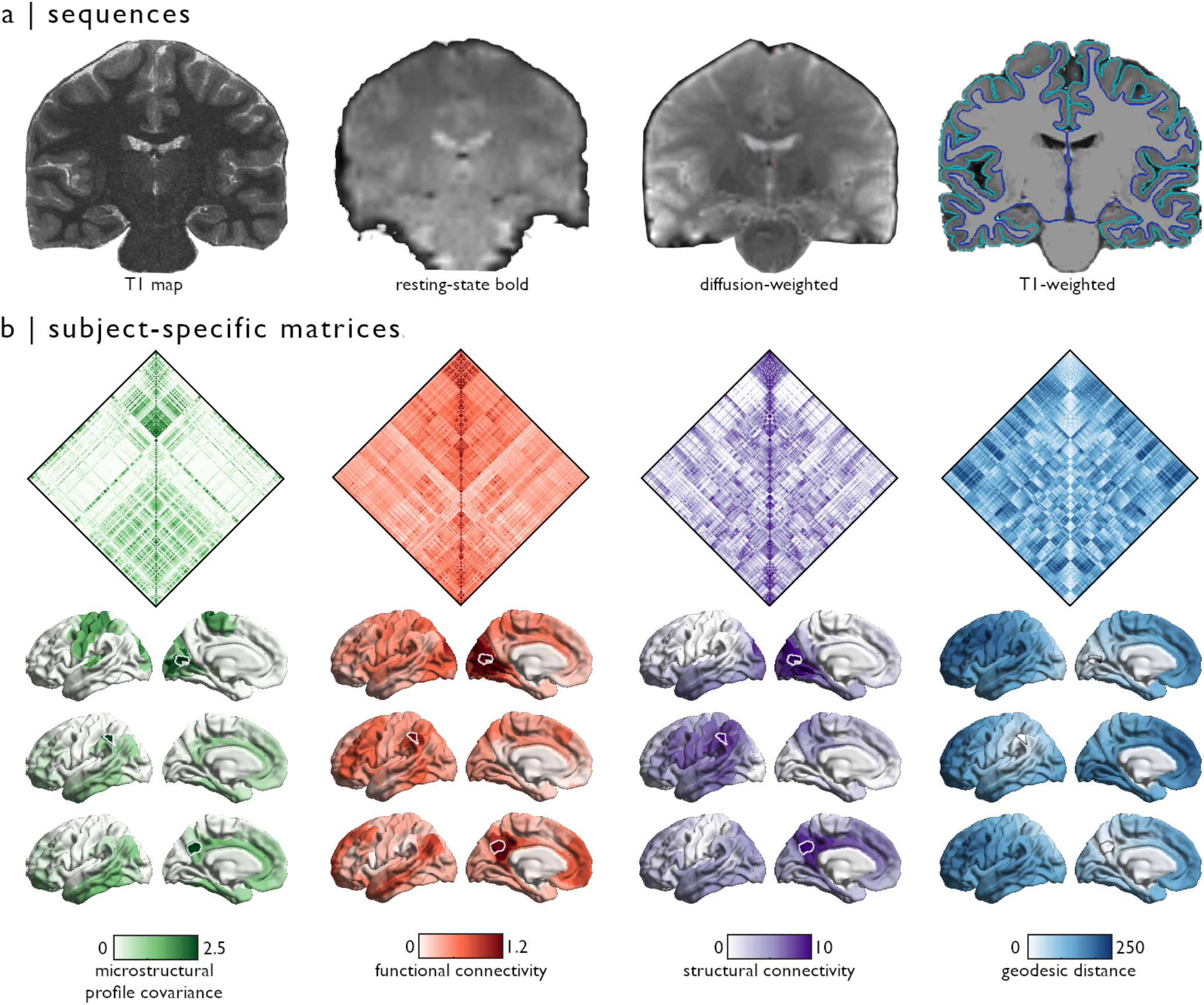
Overview of MICA-MICs dataset. **(a)** Sequences provided in the MICA-MICs dataset release include quantitative T1 relaxometry, a multiband accelerated resting-state functional scan, multiband, multi-shell diffusion-weighted imaging, and two structural T1w scans. Pial and white matter surface segmentations are superimposed on a coronal slice of the T1w image generated by FreeSurfer combining both input T1w scans. **(b)** Group-averaged matrices (only left hemisphere parcels shown - top panel) and connection weights from three outlined seeds, selected to represent a diverse set of network communities (bottom panel). Microstructural profile covariance (MPC), functional connectivity (FC), and geodesic distance (GD) matrices were averaged across participants. Group-level structural connectivity (SC) was computed using distance-dependent thresholding to preserve the distribution of within- and between-hemisphere connections lengths in individual subjects ^79^. Prior to averaging, subject-level SC matrices were log-transformed to reduce connectivity strength variance. All features are projected to the fsaverage5 midsurface from the Schaefer-400 atlas.

Two T1w scans with identical parameters were acquired with a 3D magnetization-prepared rapid gradient-echo sequence (MP-RAGE; 0.8mm isotropic voxels, matrix=320×320, 224 sagittal slices, TR=2300ms, TE=3.14ms, TI=900ms, flip angle=9°, iPAT=2, partial Fourier=6/8). Both T1w scans were visually inspected to ensure minimal head motion before they were submitted to further processing. qT1 relaxometry data were acquired using a 3D-MP2RAGE sequence (0.8mm isotropic voxels, 240 sagittal slices, TR=5000ms, TE=2.9ms, TI 1=940ms, T1 2=2830ms, flip angle 1=4°, flip angle 2=5°, iPAT=3, bandwidth=270 Hz/px, echo spacing=7.2ms, partial Fourier=6/8). We combined two inversion images for qT1 mapping in order to minimise sensitivity to B1 inhomogeneities and optimize intra- and inter-subject reliability ^52,53^. A 2D spin-echo echo-planar imaging sequence with multi-band acceleration was used to obtain DWI data, consisting of three shells with b-values 300, 700, and 2000s/mm^2^ and 10, 40, and 90 diffusion weighting directions per shell, respectively (1.6mm isotropic voxels, TR=3500ms, TE=64.40ms, flip angle=90°, refocusing flip angle=180°, FOV=224×224 mm^2^, slice thickness=1.6mm, multi-band factor=3, echo spacing=0.76ms). b0 images acquired in reverse phase encoding direction are also provided for distortion correction of DWI scans. One 7 min rs-fMRI scan was acquired using multiband accelerated 2D-BOLD echo-planar imaging (3mm isotropic voxels, TR=600ms, TE=30ms, flip angle=52°, FOV=240×240mm^2^, slice thickness=3mm, mb factor=6, echo spacing=0.54ms). Participants were instructed to keep their eyes open, look at a fixation cross, and not fall asleep. We also include two spin-echo images with reverse phase encoding for distortion correction of the rs-fMRI scans (3mm isotropic voxels, TR=4029 ms, TE=48ms, flip angle=90°, FOV=240×240mm^2^, slice thickness=3mm, echo spacing=0.54 ms, phase encoding=AP/PA, bandwidth= 2084 Hz/Px). A complete list of acquisition parameters is provided in the detailed imaging protocol available alongside this data release.

### MRI data pre-processing

Raw DICOMS were sorted by sequence, converted to NIfTI format using dcm2niix (v1.0.20200427; https://github.com/rordenlab/dcm2niix) ^54^, renamed, and assigned to their respective subject-specific directories according to BIDS ^51^. Agreement between the resulting data structure and BIDS standards was ascertained using the BIDS-validator (v1.5.10; DOI: <u>10.5281/zenodo.3762221</u>) ^55^. All further processing was performed via micapipe, an openly accessible processing pipeline for multimodal MRI data (https://micapipe.readthedocs.io/), and BrainSpace, a toolbox for macroscale gradient mapping (https://brainspace.readthedocs.io/) ^56^.

#### T1w pre-processing

Native structural images were anonymized and de-identified by defacing all structural volumes using custom scripts (https://github.com/MICA-LAB/micapipe/; *micapipe_anonymize*). Note that processing derivatives were generated from non-anonymized images. Structural processing was carried out using several software packages, including tools from AFNI, FSL, and ANTs ^57^. Each T1w scan was deobliqued and reoriented to standard neuroscience orientation (LPI: left to right, posterior to anterior, and inferior to superior). Both scans were then linearly co-registered and averaged, automatically corrected for intensity nonuniformity ^58^, and intensity normalized. Resulting images were skull-stripped, and subcortical structures were segmented using FSL FIRST ^59^. Cortical surface segmentations were generated from native T1w scans using FreeSurfer 6.0 ^60–62^.

#### qT1 pre-processing

Native qT1 scans were anonymized and de-identified by defacing. For pre-processing, a series of equivolumetric surfaces were first constructed for each participant between pial and white matter boundaries. These surfaces were used for systematic sampling of qT1 image intensities, to compute individual microstructural profile similarity matrices ^6,7^ (see next section). Here, qT1 images were co-registered to native FreeSurfer space of each participant using boundary-based registration ^63^.

#### DWIpre-processing

DWI data were pre-processed using MRtrix ^64,65^. DWI data was denoised ^66,67^, underwent b0 intensity normalization ^58^, and were corrected for susceptibility distortion, head motion, and eddy currents using a reverse phase encoding from two b=0s/mm^2^ volumes. Required anatomical features for tractography processing (*e.g*., tissue type segmentations, parcellations) were non-linearly co-registered to native DWI space using the deformable SyN approach implemented in ANTs ^68^. Diffusion processing was performed in native DWI space.

#### rs-fMRIpre-processing

rs-fMRI images were pre-processed using AFNI ^69^ and FSL ^59^. The first five volumes were discarded to ensure magnetic field saturation. Images were reoriented, as well as motion and distortion corrected. Motion correction was performed by registering all timepoint volumes to the mean volume, while distortion correction leveraged main phase and reverse phase field maps acquired alongside rs-fMRI scans. Nuisance variable signal was removed using an ICA-FIX ^70^ classifier trained in-house on a subset of 30 participants (15 healthy controls, 15 drug-resistant epilepsy patients) and by performing spike regression using motion outlier outputs provided by FSL. Volumetric timeseries were averaged for registration to native FreeSurfer space using boundary-based registration ^63^, and mapped to individual surface models using trilinear interpolation. Nativesurface cortical timeseries underwent spatial smoothing once mapped to each individual’s cortical surface models (Gaussian kernel, FWHM=10mm) ^71,72^, and were subsequently averaged within nodes defined by several parcellation schemes (see below). Parcellated subcortical timeseries are also provided in this release and were appended before cortical timeseries. Subject-specific subcortical parcellations were non-linearly registered to each individual’s native fMRI space using the deformable SyN approach implemented in ANTs ^68^.

### Generating individual and group-level connectome matrices

The following sections describe the construction of feature matrices, derived from each imaging sequence included in this data release (**Figure 1B**). Cortical connectomes are provided according to anatomical ^20^, intrinsic functional ^24^, and multimodal parcellation schemes ^26^ at different resolutions, for a total of 18 distinct cortical parcellations. Anatomical atlases available in this dataset include Desikan-Killiany (aparc) ^20^ and Destrieux (aparc.a2009s) ^21^ parcellations provided by FreeSurfer, as well as an in vivo approximation of the cytoarchitectonic parcellation studies of Von Economo and Koskinas ^73^. We additionally include similarly sized subparcellations, constrained within the boundaries of the Desikan-Killany atlas ^20^, providing matrices with 100 to 400 cortical parcels following major sulco-gyral landmarks. Parcellations based on intrinsic functional activity (Schaefer atlases based on 7-network parcellation) are also included in this release according to a wide range of resolutions (100-1000 nodes) ^24^. Lastly, we also provide connectome matrices generated from a multimodal atlas with 360 nodes derived from the Human Connectome Project dataset, known as the Glasser parcellation ^26^. All atlases are available on Conte69 ^56^ and fsaverage5 surface templates (see *parcellations* in https://github.com/MICA-LAB/micapipe), and were resampled to each participant’s native surface to generate modality- and subject-specific matrices. In addition, structural and functional connectome matrices include data for each subcortical structure (nucleus accumbens, amygdala, caudate nucleus, pallidum, putamen, and thalamus) and the hippocampus appended before entries for cortical parcels (see **Usage notes**).

#### Geodesic distance (GD)

We computed individual GD matrices along each participant’s native cortical midsurface using workbench tools ^71,72^. First, a centroid vertex was defined for each cortical parcel by identifying the vertex with the shortest summed Euclidean distance from all other vertices within its assigned parcel. The GD between centroid vertices and all other vertices on the native midsurface mesh was computed using Dijkstra’s algorithm. Notably, this implementation computes distances not only across vertices sharing a direct connection, but also across pairs of triangles which share an edge to mitigate the impact of mesh configuration on calculated distances. Vertex-wise GD values were averaged within parcels.

#### Microstructural profile covariance (MPC)

We generated 14 equivolumetric intracortical surfaces ^74^ to sample qT1 intensities across cortical depths, yielding distinct intensity profiles reflecting the intracortical microstructural composition at each cortical vertex. This number of surfaces was selected based on recent stability analyses of resulting MPC matrices ^6,7^. Data sampled from surfaces closest to the pial and white matter boundaries were discarded to mitigate partial volume effects. Vertex-wise intensity profiles were averaged within parcels. Nodal microstructural profiles were cross-correlated across the cortical mantle using partial correlations while controlling for the average cortex-wide intensity profile, and log-transformed ^6,7^. Left and right medial walls, as well as non-cortical areas such as corpus callosum and peri-callosal regions of the Desikan-Killiany and Destrieux parcellations were excluded when averaging cortex-wide intensity profiles. Resulting matrices thus represented participant-specific similarity matrices in myelin proxies across the cortex.

#### Diffusion MRI tractography derived structural connectivity (SC)

Structural connectomes were generated with MRtrix from pre-processed DWI data ^64,65^. We performed anatomically-constrained tractography using tissue types (cortical and subcortical grey matter, white matter, cerebrospinal fluid) segmented from each participant’s pre-processed T1w images registered to native DWI space ^75^. We estimated multi-shell and multi-tissue response functions ^76^ and performed constrained spherical-deconvolution and intensity normalization ^77^. We generated a tractogram with 40 million streamlines (maximum tract length=250; fractional anisotropy cutoff=0.06). We applied spherical deconvolution informed filtering of tractograms (SIFT2) to reconstruct whole brain streamlines weighted by cross-sectional multipliers ^78^. The reconstructed cross-section streamlines were mapped to each parcellation scheme (cortical and subcortical), which were also warped to DWI space. The connection weights between nodes were defined as the weighted streamline count.

#### Functional connectivity (FC)

Individual rs-fMRI timeseries mapped to subject-specific surface models were averaged within cortical parcels. The subcortical parcellation was warped to each subject’s native fMRI volume space and used to average timeseries within each node. Individual functional connectomes were generated by cross-correlating all nodal timeseries. For analyses presented in this paper, correlation values subsequently underwent Fisher-R-to-Z transformations. However, all FC matrices are provided as raw correlation matrices in the released data.

## Data records

All files are organized according to the Brain Imaging Directory Structure (BIDS) ^51^ and are hosted on the Canadian Open Neuroscience Platform’s data portal (CONP; https://portal.conp.ca/dataset?id=projects/mica-mics).

### Native space data

Native space data and corresponding .json files are contained in the branch */rawdata/sub-HC#/ses-01* of the directory structure (**Figure 2A**). For each subject (*/sub-HC#/ses-01*), the */anat* subdirectory includes several NIfTI files containing native space T1w and qT1 images. T1w scans are named according to acquisition order, denoted by *run-#*. For unprocessed qT1 images, we provide results of each inversion time parameter (denoted by *inv-1* and *inv-2*), T1 mapping based on the combination of both inversion time images (*T1-map*), as well as MP2RAGE-derived synthetic T1w images (*uni*). Removal of facial features by masking was the only change applied to these images (see *MRI data pre-processing*).

**Figure 2.**
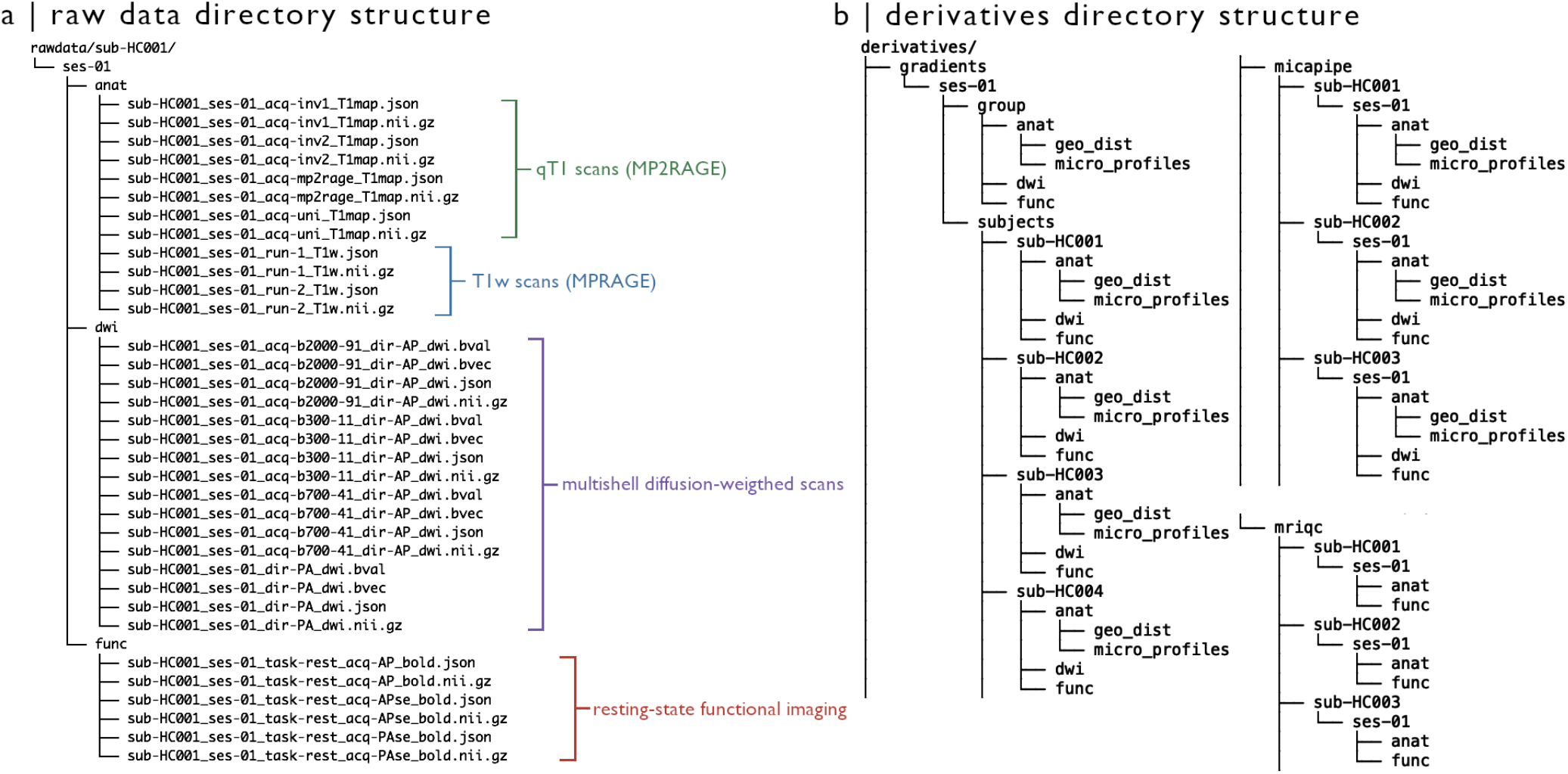
Directory structure of MICA-MICs dataset. **(A)** Anonymized data with no additional processing are provided in the *rawdata* branch of the directory structure, and includes qT1, T1w, diffusion-weighted, and resting-state functional imaging data. **(B)** Processing derivatives are organized according to their associated pipelines. Group and subject-level gradients (*/derivatives/gradients*) were derived from averaged and individual connectivity matrices computed from several parcellation schemes using micapipe (*/derivatives/micapipe*). Matrices and gradients are organized into modality-specific directories for structural (*/anat/micro_profiles* for MPC, */anat/geo_dist* for geodesic distance), functional (*/func*), and diffusion-weighted (*dwi*) imaging. We additionally provide detailed image quality reports for T1w and rsfMRI raw data generated using MRIQC ^80^.

Subject-specific DWI files can be found in the /*rawdata/sub-HC*#/*ses-01*/*dwi* subdirectory. Gradient direction, diffusion weighting, DWI volumes, and .json sidecar files are associated with each shell, indicated by its corresponding b-value and number of diffusion directions in the filename (*e.g*., “*sub-HC#_ses-01_acq-b#_dir-AP_dwi.jsorn*”). b0 images are denoted by their inverse phase encoding direction (PA; *i.e*., “*sub-HC#_ses-01_dir-PA_dwi.json*”).

The rs-fMRI scans as well as associated spin-echo images used for distortion correction are located in the /*rawdata/sub-HC#/ses-01/func* subdirectory. Functional timeseries include 700 timepoints, with the exception of subject numbers equal to or preceding *sub-HC004* who underwent slightly longer acquisition (800 timepoints). Phase encoding direction of spin-echo images are indicated in the filename (*i.e*., *APse* – anterior-posterior – or *PAse* – posterior-anterior. The string “*se*” following phase-encoding direction in the filename indicates a spinecho image later used for distortion correction).

### Processed data

Processed data included in this release are stored in the *derivatives* subdirectory associated with their processing pipeline (**Figure 2B**). Quality control reports of raw structural and functional data are provided in *derivatives*/*mriqc*/. Modality-specific matrices of varying granularity (70-1000 nodes) were generated using micapipe, and are stored in their respective subdirectory (*e.g*., all functional connectomes can be found in *derivatives*/*micapipe*/*sub-HC*#/*ses-01*/*func*/). We also provide group- and subject-level gradients generated from each matrix, stored in *derivatives*/*gradients*/*ses-01*/ (see **Technical validation and derivative metrics**).

#### Structural processing

Surface-mapped processing derivatives of structural scans are provided in /*derivatives*/*micapipe*/*sub-HC*#/*ses-01*/*anat*. These features are organized in two distinct subdirectories. First, MPC matrices generated from processed qT1 scans are stored in the /*micro_profiles* subdirectory and are identified by the parcellation scheme from which they were computed (*e.g*., “*sub-HC*#_*ses-01*_*space-fsnative*_*atlas-schaefer100_desc-mpc.txt*”). GD matrices for each cortical parcellation scheme are included in the /*geo_dist* subdirectory (*e.g*., “*sub-HC#_ses-01_space-fsnative_atlas-schaefer100_desc-gd.txt*”). As described in a previous section, individual geodesic distance matrices were computed along each participant’s native midsurface using workbench ^71,72^.

#### DWIprocessing

Processing derivatives of DWI scans are provided in /*derivatives*/*micapipe*/*sub-HC#*/*ses-01*/*dwi*. Structural connectomes (*e.g*., “*sub-HC#_ses-01_space-dwinative_atlas-schaefer100_desc-sc.txt*”) and associated edge lengths (*e.g*., “*sub-HC#_ses-01_space-dwinative_atlas-schaefer100_desc-edgeLength.txt*”) are provided for each parcellation.

#### rs-fMRIprocessing

Fully processed connectomes (*i.e*., after removal of nuisance variable signal using ICA-FIX ^70^, mapping to native cortical surface, spatial smoothing, and regression of motion spikes) are provided in /*derivatives*/*micapipe*/*sub-HC#*/*ses-01*/*func* (*e.g*., “*sub-HC#_ses-01_space-fsnative_atlas-schaefer100_desc-fc.txt*”). Functional connectomes were computed from native-surface mapped timeseries for congruency across data modalities, as both GD and MPC matrices are generated from data mapped to native cortical surface models.

#### Quality control

Reports of image quality metrics computed by MRIQC v0.15.2 (https://github.com/poldracklab/mriqc/) ^80^ are included in the /*mriqc* branch of MICA-MICs processing derivatives. For each subject, /*mriqc* directories contain /*anat* and /*func* subdirectories, which include image quality metric reports for T1w and resting-state functional scans in .html and json formats. These reports provide a number of metrics evaluating the quality of the input data, including estimates of motion, signal-to-noise, and intensity non-uniformities ^80^.

## Technical validation and derivative metrics

### Quality control procedures

#### Cortical surface segmentations

Surface extractions were visually inspected by three authors (JR, AJL, CP) and corrected for any segmentation errors with the placement of control points and manual edits.

#### Image quality metrics

The consistency of T1w scan quality was assessed using contrast-to-noise estimates computed in MRIQC ^80^ (**Figure 3A**). This metric provides a measure of separability of grey and white matter distributions for a given T1w image ^80,81^, with higher values indicating better image quality. For DWI scans, movement was quantified in each shell using MRtrix and FSL eddy ^82^ (**Figure 3B**). For rs-fMRI, framewise displacement (FD) was estimated using FSL’s motion outlier detection tool. We also explored temporal signal-to-noise (tSNR) ratio, calculated for each participant by dividing surface-mapped mean timeseries by their standard deviation. Motion and distortion corrected timeseries were used to calculate tSNR across the cortex for each participant (*i.e*., before high-pass filtering and nuisance signal regression using ICA-FIX). Vertex-wise tSNR values were averaged within parcels to aggregate values across subjects (**Figure 3C**).

**Figure 3.**
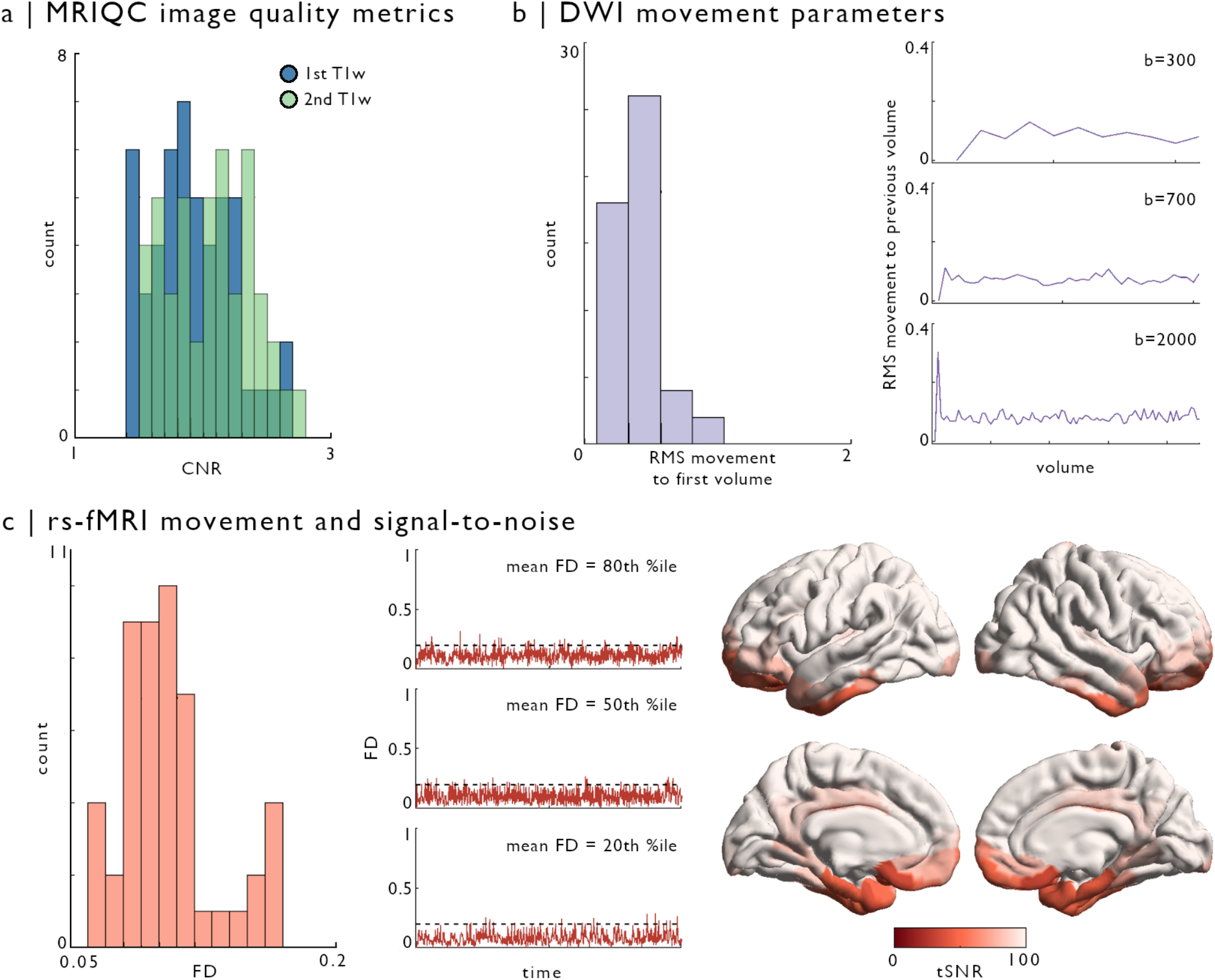
Image quality metrics across sequences. **(A)** Contrast-to-noise (CNR), estimated with MRIQC ^80^, showed no outliers in either T1w scan (first scan in blue, second scan in green). **(B)** Motion parameters of diffusion-weighted images were obtained from FSL eddy ^82^. The histogram illustrates root mean squared (RMS) voxel-wise displacement relative to the first volume across all shells. Line plots show RMS displacement in each volume relative to the previous volume. **(C)** Framewise displacement (FD) of resting-state functional scans was obtained using FSL motion outliers, reflecting the average of rotation and translation parameter differences at each volume ^83^. The histogram shows subjects-wise average FD across volumes. Line plots show FD across resting-state acquisitions for three participants, with respectively 20^th^, 50^th^, and 80^th^ percentile average FD across our sample. Dashed line indicates 0.2 mean FD threshold used for exclusion of participants with excessive motion. Vertex-wise temporal signal-to-noise (tSNR) was calculated on the native surface of each participant. Computed tSNR values were averaged within a 400-node functional parcellation (Schaefer-400) and averaged across individuals.

### Estimation of cortical gradients from MPC, FC, SC, and GD matrices

In this section, we demonstrate how group and individual-level gradients can be derived from each data modality provided in MICA-MICs. Using the BrainSpace toolbox (http://brainspace.readthedocs.io) ^56^, we identified gradients from MPC, FC, SC, and GD matrices. We constructed group-level gradients by averaging all cortical entries of subject-level matrices constructed from the Schaefer-400 atlas. MPC, FC, and GD matrices were computed by cross-subject averaging, and results were thresholded row-wise to retain the top 10% edges, as in previous work ^6,14,30,34^. Group-level structural connectivity (SC) was computed using distance-dependent thresholding to preserve the distribution of within- and between-hemisphere connection lengths in individual subjects ^79^. Prior to averaging, subject-level SC matrices were log-transformed to reduce connectivity strength variance. Group-average SC matrices were thresholded to only retain positive edges. No further thresholding was applied given the sparsity of SC matrices relative to other modalities.

Normalized angle affinity matrices, capturing inter-regional similarity of microstructural, connectivity, and distance patterns, were computed from each modality-specific matrix (**Figure 4A**, top). Left and right hemispheres were analysed separately for SC data, given limitations of diffusion tractography in mapping inter-hemispheric fibres. Hemispheres were also analysed separately for GD gradients, as the surface-based measure of geodesic distance used here is computed on distinct hemisphere surface spheres. Data from both hemispheres were used to generate affinity matrices from MPC and FC features. We applied diffusion map embedding, a non-linear dimensionality reduction technique ^14,56,84^, to each affinity matrix to identify eigenvectors (or gradients) describing inter-regional variability in each feature in descending order for each modality (**Figure 4A**, middle). Resulting gradients were visualized on cortical surfaces, revealing distinct patterns for each feature (**Figure 4A**, bottom). For instance, the first MPC gradient (G1) derived from myelin-sensitive qT1 recapitulated a sensory-fugal axis ^45,46^ ordering nodes from sensorimotor to paralimbic cortices ^6^. In contrast, the principal FC and SC gradients primarily distinguished visual and sensorimotor cortices. The second gradient of FC, explaining a similar amount of variance to FC-G1, was anchored in unimodal sensory systems and the higher-order default mode network ^14^. Gradients of geodesic distance highlighted the longest distance axes across the cortical surface mesh, specifically evolving along anterior to posterior (G1) and mesial/inferior to lateral/superior (G2) directions.

**Figure 4.**
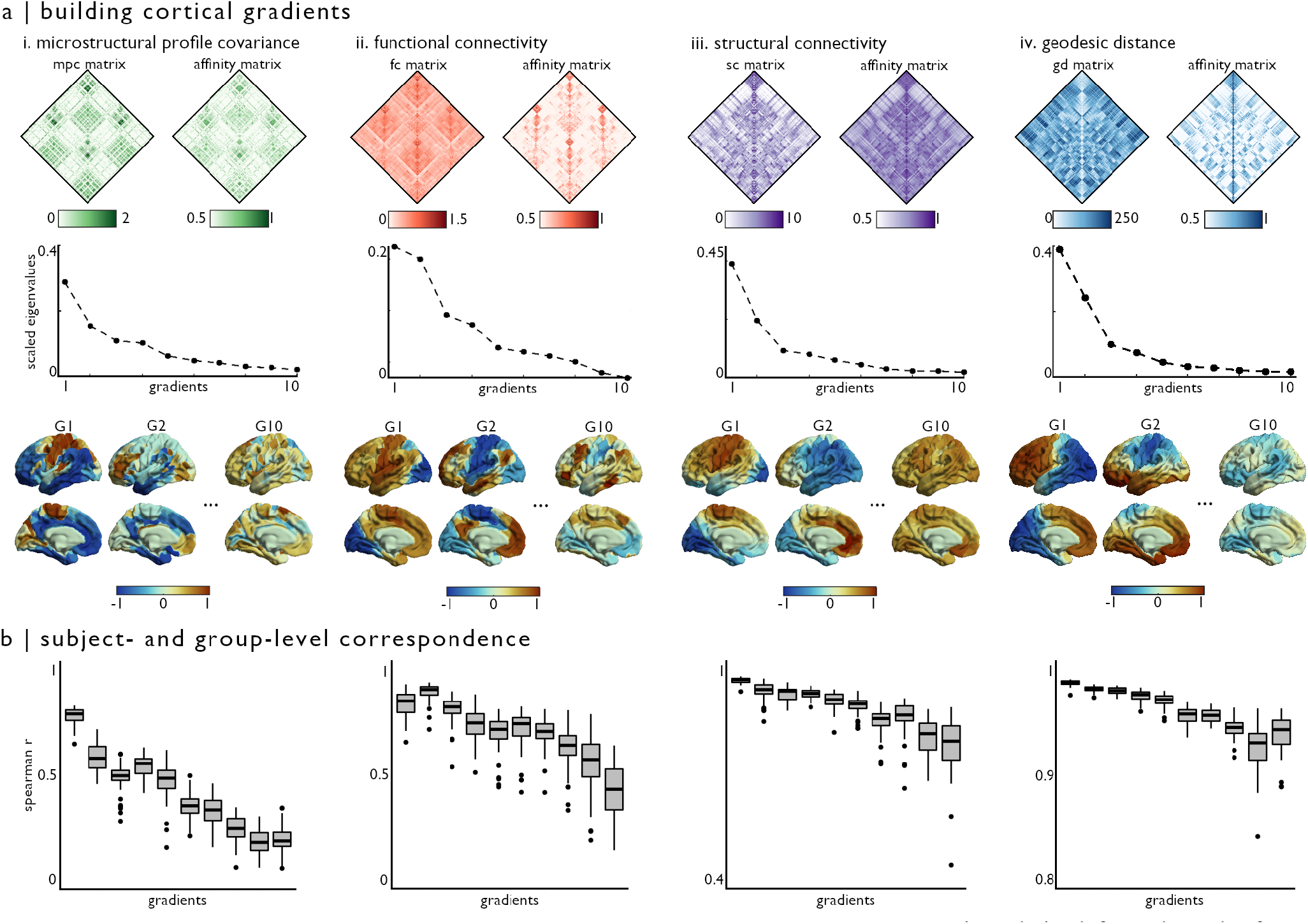
Deriving smooth microstructural, connectivity, and distance gradients. **(a)** Matrices derived from the Schaefer-400 parcellation describing i) microstructural similarity, ii) functional connectivity, iii) structural connectivity, and iv) spatial proximity were thresholded, and transformed into affinity matrices using a normalized angle kernel (top row). Only left hemisphere data is shown, although data from both hemispheres was included in MPC and FC analyses. We then applied diffusion map embedding, a non-linear dimensionality reduction technique, to each affinity matrix to derive gradients describing inter-regional variability in each feature in descending order (middle row). A subset of resulting gradients is projected onto the cortical surface for each modality (bottom row). **(b)** We assessed reproducibility of group-level gradient patterns at the individual-participant level using Spearman correlations. We generated gradients for each modality, in each participant, and aligned resulting eigenvectors to corresponding group-level gradient data. Box plots show variations in Spearman r-values across participants, for the first 10 gradients in each modality (presented in the same order as panel a). Note change in y-axis scale in SC and GD box plots.

We next assess the reproducibility of group-average gradients in individual participants. Subject-level gradients were generated following the same procedure as previously described group-level analyses. Resulting subject-level gradients were aligned with group-level template gradients generated from the 49 other participants using Procrustes alignment ^56^. This procedure (*i.e*., excluding a single participant from the template used for alignment) ensured that resulting correlations were not spuriously increased by correlating single-subject data present in both sets. Aligned subject-level gradients were correlated with their corresponding gradient in the group-level data (**Figure 4B**). A similar pattern was seen across all modalities, with decreasing individual-level replicability in gradients explaining less variance within each feature. Indeed, G1 was highly reproducible in all participants across all modalities (r mean±SD; MPC 0.785±0.041; FC 0.839±0.065; SC 0.973±0.008; GD 0.989±0.003), but correlations between individual subject data and group-level template gradients were lower for gradients explaining less variance (*e.g*., G10; MPC 0.193±0.064; FC 0.416±0.127; SC 0.785±0.083; GD 0.940±0.019).

All subject-level gradients provided in this release were aligned to the full group template, and are provided for each modality and parcellation scheme. As such, all individual-subject gradients are aligned to an identical template. These files are included in their respective /*derivatives* subdirectories. For instance, all FC gradients for a given participant can be found in the /*derivatives*/*gradients*/*ses-01*/*subjects*/*sub-HC*# subdirectory (*e.g*., “*sub-HC#_ses-01_space-fsnative_atlas-schaefer100_desc-fcGradient.txt*” for FC gradients). Gradients generated from the averaged full sample data can also be accessed within their respective /*derivatives/gradients* directories (*e.g*., /*derivatives*/*gradients*/*ses-01*/*group*/*func* for FC gradients).

## Usage notes

### Data hosting

MICA-MICs is made openly available via the CONP portal (https://portal.conp.ca/dataset?id=projects/mica-mics).

### Matrix ordering

Rows and columns of GD and MPC matrices follow the order defined by annotation labels associated with their parcellation (see *parcellations* in https://github.com/MICA-LAB/micapipe), including unique entries for the left and right medial walls. For example, row and column entries of the Schaefer-100 matrices are ordered according to: Left hemisphere cortical parcels (1 medial wall followed by 50 cortical regions), and right hemisphere cortical parcels (1 medial wall followed by 50 cortical regions). FC and SC matrices follow the same ordering, although entries for subcortical structures are appended before cortical parcels. As such, row and column entries of the Schaefer-100 FC and SC matrices are ordered according to: Subcortical structures and hippocampus (7 left, 7 right), left hemisphere cortical parcels (1 medial wall followed by 50 cortical regions), and right hemisphere cortical parcels (1 medial wall followed by 50 cortical regions). The ordering of all parcels and their corresponding label in each volumetric parcellation are documented in lookup tables provided with our analysis pipeline.

### Gradient data

Nodes excluded from group- and individual-level gradient analyses are indicated by a value of *Inf* in the corresponding node index. These data points may correspond to non-cortical nodes (*e.g*., medial wall, callosal or peri-callosal areas) or to nodes with no connections to other areas. This second case occasionally occurred in higher-resolution (>500 nodes) SC matrices of individual subjects.

## Code availability statement

All processing pipeline scripts are openly available. Code used to generate pre-processed outputs can be accessed via GitHub (https://github.com/MICA-LAB/micapipe). Documentation for the processing pipeline, including usage and detailed processing steps, can also be accessed via ReadTheDocs (https://micapipe.readthedocs.io).

## Author contributions

Conception, design, and manuscript preparation: JR, BCB; Participant recruitment: JR, ST; Data acquisition: QL, AJL, JR, ST; Processing pipeline: PH, SL, CP, BP, JR, RRC, RV; Data processing and quality control: SL, QL, JR, RRC, ST, RV. All authors provided feedback and approved the final manuscript.

## Acknowledgements

The authors wish to thank all participants who took part in this study. We also thank David Costa, Ronald Lopez, and Louise Marcotte for their assistance in data collection, as well as Ilana Leppert and Michael Ferreira for their guidance in designing the scanning protocol.

